# Engineered subtilisin protease degrades active KRAS in cancer cells, leading to differential cell targeting

**DOI:** 10.64898/2026.03.04.709402

**Authors:** M.E. Goldstein, B. Chu, K.J. Carillo, J. Orban, E.A. Toth, T.R. Fuerst

## Abstract

Controlling aberrant RAS signaling has been the subject of intensive efforts aimed at developing specific RAS inhibitors, small molecules that promote RAS degradation, and monobodies that inhibit RAS activity. Direct proteolytic degradation of RAS by site-specific proteases has received considerably less attention. A naturally-occurring protease from *Vibrio vulnificus* toxin cleaves all RAS isoforms at switch I and attenuates RAS signaling in cell models and patient-derived xenografts, thus demonstrating the potential of this approach. We previously designed a RAS-specific protease, called RASProtease (or RASp), that site-specifically cleaves RAS at switch II. Attacking switch II leverages an order to disorder transition that this region undertakes upon conversion to the active form that predominates in cancer. Switch II participates in an allosteric network that controls KRAS oncogenicity, making it a promising target for proteolytic cleavage that modulates RAS signaling. Preferential targeting of active RAS could be particularly useful for studying RAS signaling networks as well as having potential therapeutic value. Here we examined the effects of RASp cleavage on downstream signaling and cell viability in the MIA PaCa-2 cancer cell model, which harbors homozygous KRAS G12C and is KRAS-dependent for growth and survival. We found that cleavage of KRAS G12C coincided with a decrease in MEK-ERK signaling and resulted in extensive MIA PaCa-2 cell death 24 hours after induction of RASp expression. This level of cell death far exceeded that of control HEK 293T cells under the same conditions, underscoring the vulnerability of this cancer cell model to KRAS G12C elimination.

## Introduction

The rat sarcoma (*RAS*) gene family of small-GTPases, containing three distinct, yet overlapping in function isoforms, KRAS, NRAS, HRAS, play a critical role in regulating cell growth and proliferation (*1*). Mutations within *RAS*, result in locking these regulatory proteins in the GTP-bound activated state, and leading to uncontrolled cell proliferation and ultimately cancer (*2*). Oncogenesis occurs with the locked on-state, triggering the downstream activation of signaling effectors within the MAPK pathway, including the mitogen-activated protein kinase kinase (MEK), extracellular regulated kinase (ERK), and the mechanistic target of rapamycin (mTOR), including AKT. Within cancers, *RAS* is the most frequently mutated gene family (*2*), with mutations at position 12 resulting in the highest percentages of RAS-derived cancers (*3*).

Importantly, two regions within *RAS*, denoted switch I (amino acids 30–38) and switch II (amino acids 59–76), undergo conformational changes during the shift from the inactive, GDP-bound state to the active, GTP-bound state. It is in the GTP-bound conformation where a cascade of effectors are activated and phosphorylated, leading to downstream changes in MAPK and mTOR signaling pathways. These regions provide potential vulnerabilities to target via proteolytic degradation of these cryptic cleavage sites utilizing an engineered protease. Because the inactive state of RAS remains ordered and structurally hidden (*4, 5*), switch I and switch II provide an inherent preferential bias towards proteolytic degradation of the active state. Once deemed “undruggable” due to the duality of lacking binding pockets for potential small-molecule inhibitors and its strong affinity for its natural ligand GTP (*6*), few therapeutics are currently approved for RAS-driven cancers. To date, the only KRAS therapeutics approved for use by the FDA are not-specific for the active, GTP-bound RAS protein, and thus have heightened potential for impacting healthy RAS signaling.

Previously, we re-engineered the specificity of the *Bacillus subtilis* serine protease, subtilisin, to target switch II of RAS, which reveals an exposed unstructured region, QEEYSAM, only in the active state (*7–10*). We named this engineered subtilisin RASProtease (*8*), and for brevity we will refer to it here as RASp. Additionally, to allow tighter control of its activity, RASp was altered to require a cofactor, either imidazole (I) or nitrite (N), with the respective proteases designated RASpI and RASpN, respectively. Thus, given our initial success using RASp to cleave KRAS in HEK 293T cells, we sought to further validate the ability of RASp to prevent downstream cell signaling cascades, as well as to demonstrate the ability to degrade active RAS in more relevant cell culture models with RAS mutations specific to cancer.

Utilizing MIA-PaCa-2 (MP2) cells, an immortalized primary pancreatic cancer cell line, which harbors the KRAS G12C mutation, we define the ability of RASp to specifically degrade GTP-bound KRAS, preventing both MAPK and mTOR signaling cascades, and ultimately driving these RAS-dependent cells to a more rapid cell death compared to cells expressing wild-type (WT) KRAS. Thus, RASp, provides the foundation for re-engineering proteases towards novel and difficult-to-target cellular proteins or pathogens. Specifically, for RAS-based cancers, RASp provides a potential framework for developing a pan-RAS therapeutic.

## Results

Given our previous findings (*8*) and prior to moving into *in vitro* studies, we wanted to investigate which cofactor, imidazole or nitrite, would be better suited for use in cell models. Imidazole is an exogenous cofactor that might provide tighter regulatory control over the activation of a protease therapeutic. Nitrite, by contrast, is a naturally present cofactor, and often present at high concentrations, in cancer cells. This endogenous cofactor would be internally regulated by the nitrite concentration present in cancer cells, yet it could be more challenging to influence protease activity via supplementation of cell culture media. Thus to initially assess the biochemical activity of the two RASp variants, we compared the cleavage kinetics of RASpI and RASpN using a fluorescent peptide substrate, QEEYSAM (**Figure 1A, 1B**), where fluorophore fluorescence remains quenched until it is cleaved from the peptide. Here, RASpN cleaves the peptide substrate at a rate of approximately 1.57 s^-1^, whereas RASpI cleaves the same substrate at a rate of approximately 0.012 s^-1^. Because the cleavage rates, *k_obs_*, differ by two orders of magnitude, RASpN appears to be a more promising protease for use in cancer cell models.

**Figure 1.**
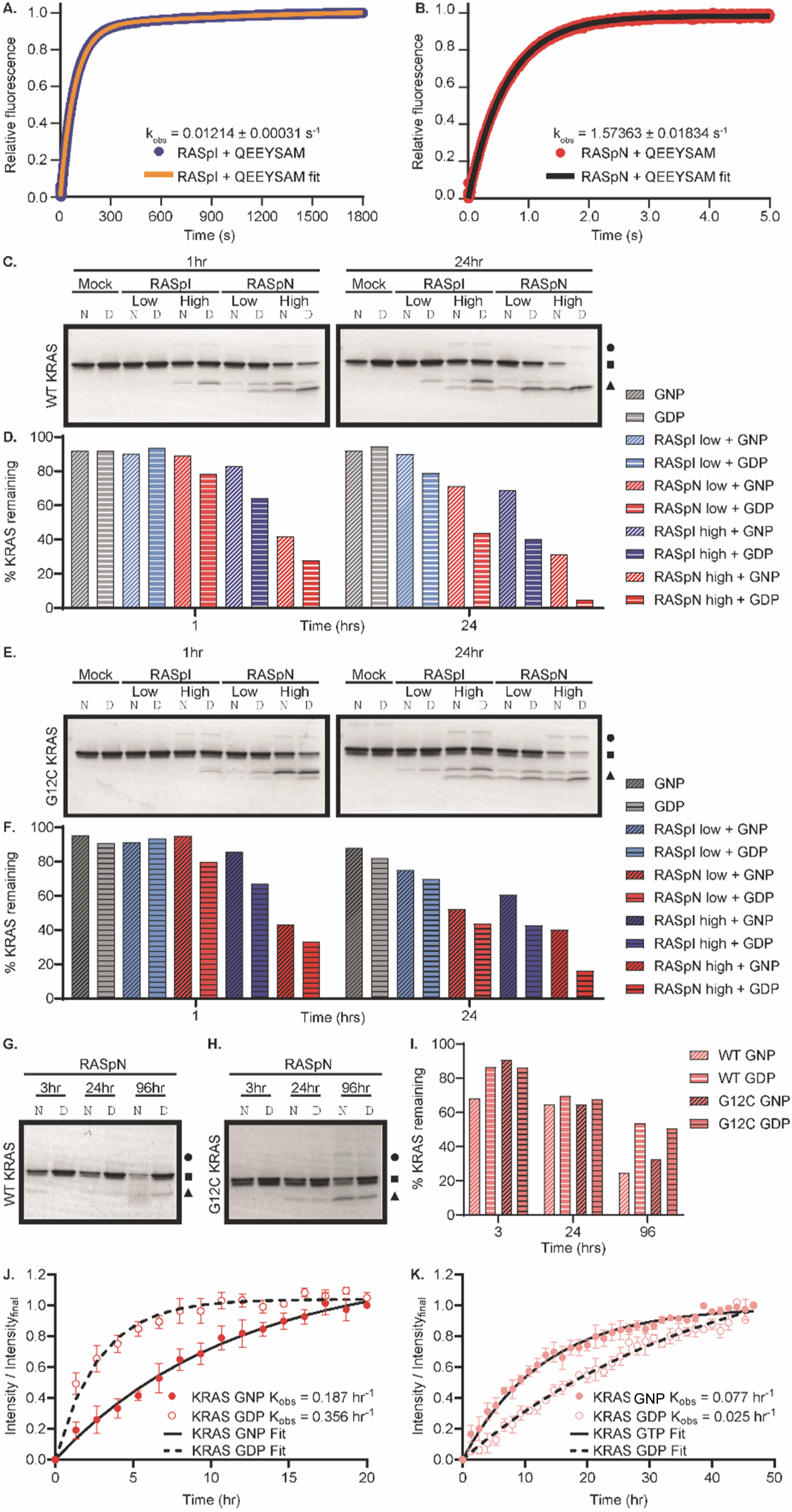
RASp cleaves purifed KRAS. (**A**) Single-turnover fluorescence kinetics of 100nM of QEEYSAM, a peptide mimetic of the switch 2 region of KRAS, after incubation with 4µM RASpI. (**B**) Single-turnover fluorescence kinetics of 100nM of QEEYSAM, a peptide mimetic of the switch 2 region of KRAS after incubation with 4µM RASpN. (**C**) Coomassie-stained gels of *in vitro* proteolysis of purified GMPPNP- (T) or GDP-bound (D) WT KRAS after incubation with stoichiometric amounts of purified RASpI or RASpN in dose-dependent manner. Proteolysis reactions were allowed to proceed for 1hr, left, 3hr, center, or 24hr, right. (**D**) Coomassie-stained gels of *in vitro* proteolysis of purified GMPPNP- (T) or GDP-bound (D) G12C-mutant KRAS after incubation with with stoichiometric amounts of purified RASpI or RASpN in dose-dependent manner. Proteolysis reactions were allowed to proceed for 1hr, left, 3hr, center, or 24hr, right. (**E**) Quantification of WT KRAS degradation in Coomassie-stained gels in (C) using FIJI. (**F**) Quantification of G12C-mutant KRAS degradation in Coomassie-stained gels in (D) using FIJI. (**G**) Coomassie-stained gels of *in vitro* proteolysis of purified GMPPNP- (T) or GDP-bound (D) WT KRAS after incubation with rate-limiting, 1:10000, amounts of purified RASpI or RASpN. Proteolysis reactions were allowed to proceed for 3hr, 24hr, or 96hr. (**H**) Coomassie-stained gels of *in vitro* proteolysis of purified GMPPNP- (T) or GDP-bound (D) G12C-mutant KRAS after incubation with rate-limiting amounts of purified RASpI or RASpN. Proteolysis reactions were allowed to proceed for 3hr, 24hr, or 96hr. (**I**) FIJI quantification of WT and G12C-mutant KRAS degradation in Coomassie-stained gels in (F,G). (**J**) RASpN kinetics after incubation with stoichiometric amounts of WT KRAS GDP, circles, and GMPPNP, triangles, by NMR. (**K**) RASpN kinetics after incubation with rate-limiting, 1:10000, amounts of WT KRAS GDP, circles, and GMPPNP, triangles, by NMR.

While these findings suggested RASpN would degrade KRAS at a faster rate than RASpI, we further advanced our understanding of RASp kinetics by incubating both RASp variants, with their respective cofactors, at both a high and low (1:10 dilution) concentrations using a stoichiometric amount of KRAS at 37 °C. To assess whether RASp can selectively prefer the active GTP-bound state over the inactive GDP-bound state, KRAS-GDP was subjected to nucleotide exchange with GMPPNP as a stable GTP analog. To better ascertain the kinetics of this cleavage reaction, we terminated the reaction at 1 hr, 3 hr, and 24 hrs and assessed KRAS cleavage by SDS-Page gel after staining with Coomassie blue (**Figure 1C**). Complementing our prior results, which used only the peptide substrate, we note more advanced KRAS degradation upon incubation with RASpN compared with RASpI. Importantly, we observe a near-complete disappearance of the WT GDP-bound KRAS band upon incubation with RASpN by 24 hrs, while the GMPPNP analog-bound KRAS remains at depleted, yet not nearly as reduced, levels. To further support these comparisons, band intensities were calculated using the FIJI software, to quantify the total amount of protein signal in each visible band (**Figure 1D**). From these transformed data, it becomes readily apparent that the low RASpN condition cleaves at a similar rate to that of high RASpI, which further indicates that RASpN is the more potent enzyme. We next assessed the ability of RASpI and RASpN to effectively degrade not only WT KRAS, but also KRAS harboring the G12C mutation. The latter is a highly prevalent mutation in a several cancers (*3*). Thus, in parallel with the experiments conducted in Figure 1C and D, we also compared the ability of RASp to degrade both GDP-bound and GMPPNP-bound KRAS G12C (**Figure 1E, 1F**). Here, using KRAS G12C protein, we also observe the ability of stoichiometric amounts of RASpN to more rapidly degrade mutant GDP-bound KRAS. However, upon repeating these proteolytic degradation assays and associated SDS-PAGE with either GTP- or GDP-bound KRAS in the presence of either WT (**Figure 1G**), or G12C mutant (**Figure 1H**), but RASpN at a concentration of 1 nM, we observe a complete switch in which substrate preference moves towards active KRAS. While temporally slower, given the 1:10000 dilution of RASpN, by 96 hpi (hours post-incubation), the GTP-bound KRAS becomes degraded more rapidly than the GDP-bound analog. These important findings were further validated by NMR analysis (**Figure 1J**), where under conditions similar to **Figure 1E**, using stoichiometric amounts of KRAS and RASpN, we observe GDP-bound KRAS to be degraded more rapidly, with a *k_obs_* value of 0.356 hr^-1^ vs 0.187 hr^-1^ for GTP-bound KRAS. However, upon incubation with 1 nM RASpN (**Figure 1K**), we note a more rapid degradation of GTP-bound KRAS (*k_obs_* ∼ 0.077 hr^-1^) compared with GDP-bound KRAS, (*k_obs_* ∼ 0.025 hr^-1^) under the same conditions. Since achieving stoichiometric equivalent amounts of RASpN to KRSAS would be difficult in cell models, and likely highly cytotoxic, the realistic concentrations of RASp will better target and degrade oncogenic KRAS.

To further our investigation into the ability of RASp to cleave KRAS in vitro, we transfected HEK 293T cells with an eGFP fusion KRAS construct, which is labeled eGFP-KRAS. At 72 hours post transfection (hpt), eGFP-KRAS-expressing cells were lysed in RIPA buffer containing protease and phosphatase inhibitors, as indicated by the experimental design cartoon (**Figure 2A**). Lysates were then incubated with either RASpI or RASpN and their respective cofactors under high, low (1:10 dilution) or mock conditions for 1 hr, 3 hr or 24 hr at 37 °C and resulting lysates were analyzed by western blot (**Figure 2B**). We observe the intact eGFP-KRAS fusion protein at 50 kDa in all eGFP-KRAS containing lanes at 1 hr but begin to observe KRAS degradation at that time point in both low and high RASpN (and the onset of cleavage in the high RASpI) conditions and the resultant degradation products upon probing with an anti-GFP polyclonal antibody, detecting free eGFP (27 kDa) and lower molecular weight protein fragments (<15 kDa). Probing with an anti-KRAS monoclonal antibody corroborates these findings via detection of a semi-stable degradation product (<15 kDa). These cleavage patterns become much more readily apparent by 24 hours post incubation (hpi), where the eGFP-KRAS fusion has become completely degraded in high RASpI and low and high RASpN conditions. Here, even the low RASpI condition has greatly reduced full-length eGFP-KRAS remaining. Importantly, when probing with anti-KRAS antibody, at 24 hpi, the high RASpN condition has completely abolished all KRAS-containing species. To strengthen these gels, blot intensities of the eGFP-KRAS fusion protein were quantified across timepoints, anti-GFP (**Figure 2C**) and anti-KRAS (**Figure 2D**); **Figure 2E** furthers these data by quantifying the decrease in eGFP-KRAS signal at 24 hpi compared with the initial 1 hr timepoint. Thus, here we validate the ability of RASp to cleave KRAS in vitro and further confirm that RASpN is the more potent enzyme.

**Figure 2.**
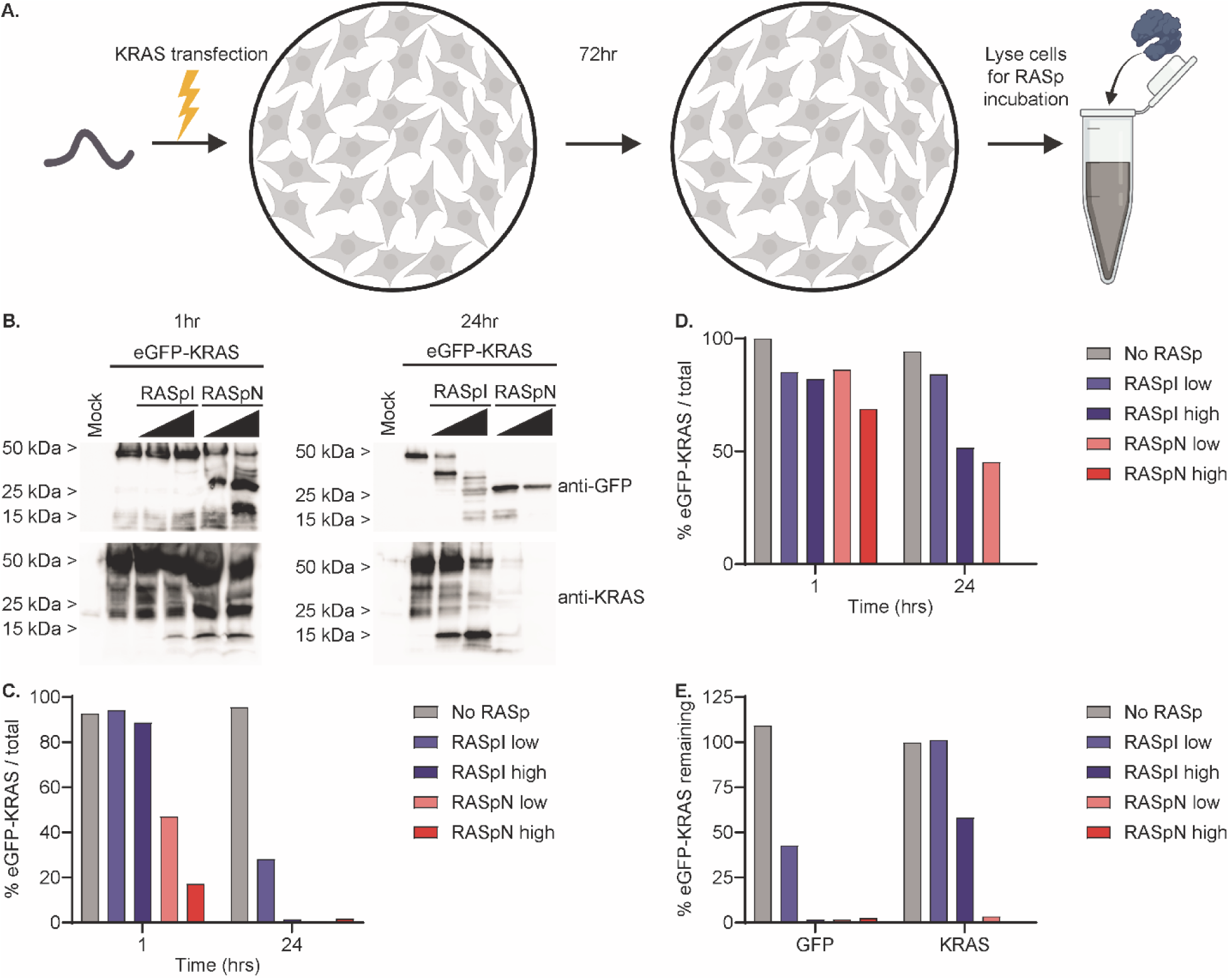
Purified RASp cleaves exogenous eGFP-KRAS expressed in HEK 293T cell lysates. (**A**) Experimental design schematic created with BioRender.com. HEK 293T cells were lysed in RIPA buffer prior to incubation with RASp. (**B**) Western blot shows cleavage of HEK 293T cell lysates expressing eGFP-KRAS after incubation with RASpI or RASpN in a dose-dependent manner for 1 hr, left, or 24 hr, right. (**C**) Quantification of KRAS expression signal by eGFP detection in (A) using FIJI. (**D**) Quantification of KRAS expression signal by KRAS detection in (A) using FIJI. (**D**) Quantification of remaining KRAS expression in (A) expressed as ratio of KRAS signal remaining at 24 hr divided by the eGFP or KRAS signal at 1 hr using FIJI.

Having identified RASpN as the more proficient enzyme using purified proteins, we sought to understand the impact of RASp expression in cells. As in Figure 2, we transfected 293T cells (**Figure 3A**) with the eGFP-KRAS construct but this time as part of a co-transfection with a FLAG-tagged, tetracycline-inducible, either RASpI, RASpN or RASp*, a RASp with a critical mutation in the active site (S221A), which renders it inactive. Expression of these RASp constructs were monitored in cells (**Figure 3B**) by detection of the FLAG tag in western blots, where we note the gradation of expression of RASp upon induction with doxycycline, from inactive, RASp*, to modestly active RASpI, to highly active RASpN. The inactive RASp* fails to auto-process its prodomain, resulting in migration at a higher molecular weight than processed RASpI and RASpN. We next looked at the effects of RASp expression on eGFP-KRAS. Upon induction of RASp expression, we observe no change in either the amount of full-length eGFP-KRAS or appearance of degradation products for RASp*, minimal reduction of full length eGFP-KRAS and background levels of cleavage products for RASpI, and a significant reduction of full length eGFP-KRAS levels and concomitant appearance of a cleavage product for RASpN. These results were further corroborated by the quantification of the band intensities using FIJI for both GFP, (**Figure 3C**), and KRAS, (**Figure 3D**). Given the robust cleavage of KRSAS by RASpN, we assessed its effects on cell health upon expression by immunofluorescence microscopy (**Figure 3E**) and counting cells remaining 72 hpt (**Figure 3F**), where we observe a near total elimination of all cells upon transfection and induction of RASpN.

**Figure 3.**
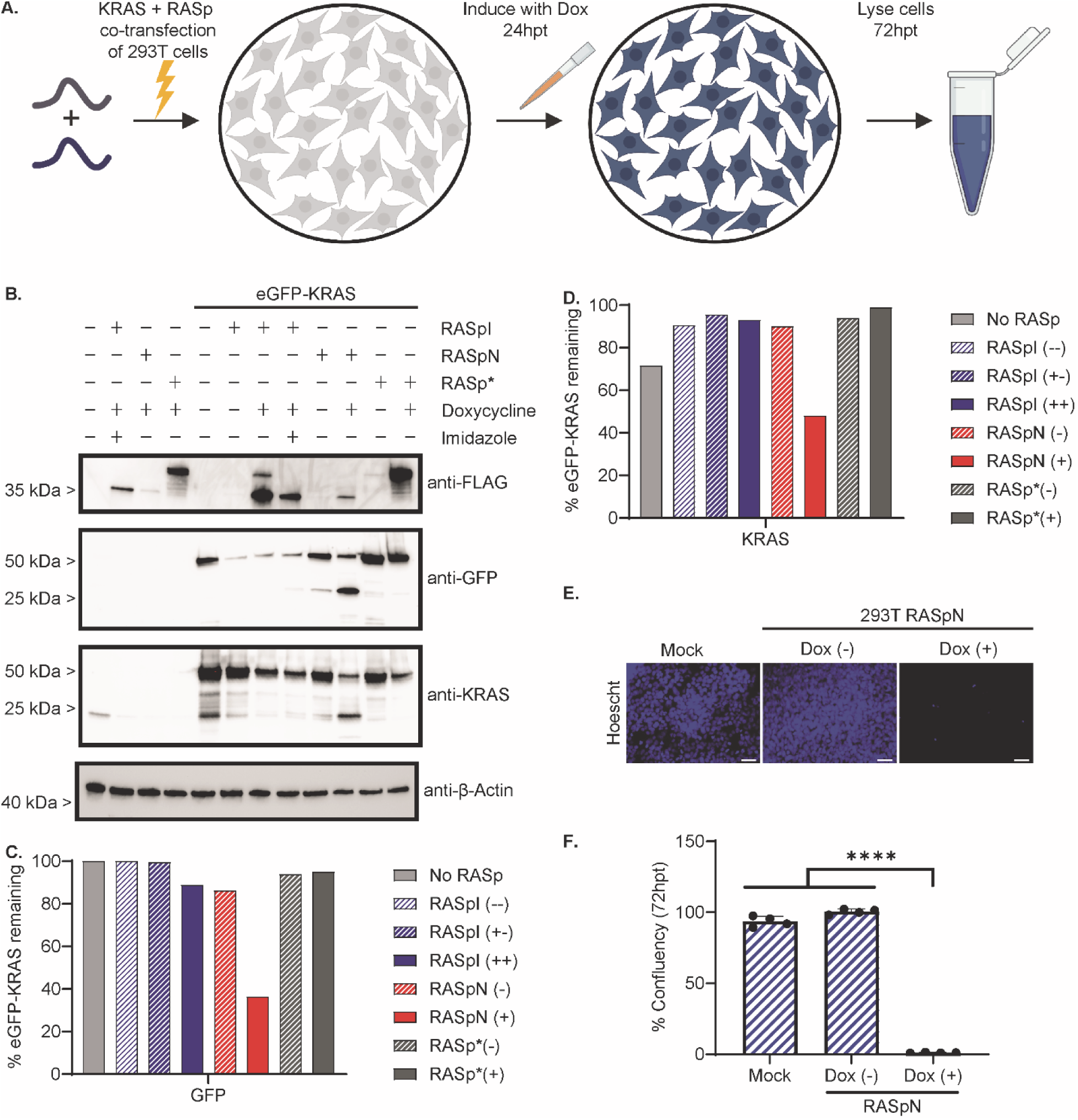
RASpN cleaves exogenous KRAS expressed by co-transfection in 293T cells. (**A**) Experimental design schematic created with BioRender.com. 293T cells were co-transfected with eGFP-KRAS and RASp. 24hpt RASp was induced with doxycycline. Cells were lysed in RIPA buffer 72hpt. (**B**) Western blot identifies cleavage of 293T cell lysates co-expressing eGFP-KRAS and RASp. (**C**) Quantification of remaining eGFP-KRAS expression signal by GFP detection in (A) using FIJI. (**D**) Quantification of remaining eGFP-KRAS expression signal by KRAS detection in (A) using FIJI. (**E**) Immunofluorescence-mediated detection of 293T cells transfected in parallel with a tetracycline-inducible RASpN or mock-transfected. RASpN-transfected cells were treated with doxycycline 24hpt, with staining performed 72hpt. Nuclei (Hoescht; blue). Scale bar = 100 µm. (**F**) Quantification of the % confluency from panel (D) using FIJI. Each bar represents mean assayed across n=4 images taken at predetermined locations within the culture. Error bars represent standard deviation across images. Statistical analysis was performed using a paired t-test (**** p <0.0001).

With these data in hand, we selected RASpN for subsequent studies in a more relevant pancreatic cancer cell line, MIA-PaCa-2 (MP2), known which harbors a homozygous KRAS G12C mutation. Similar to the experiments performed in Figure 2, we incubated MP2 cell lysates with two concentrations of purified RASpN, as indicated by the cartoon (**Figure 4A**), for 1 hr, 3 hr or 24 hr (**Figure 4B**) and observed cleavage of KRAS by western blot. After 1 hr of incubation, the lysate incubated with the high concentration of RASpN already began showing a faint lower molecular weight species indicative of proteolytic cleavage and by 24 hr of incubation, this sample was fully degraded. Cleavage of KRAS in MP2 lysates were quantified using FIJI at both 1 hr and 24 hr timepoints (**Figure 4C**) and as a percent of the original parent sample (**Figure 4D**).

**Figure 4.**
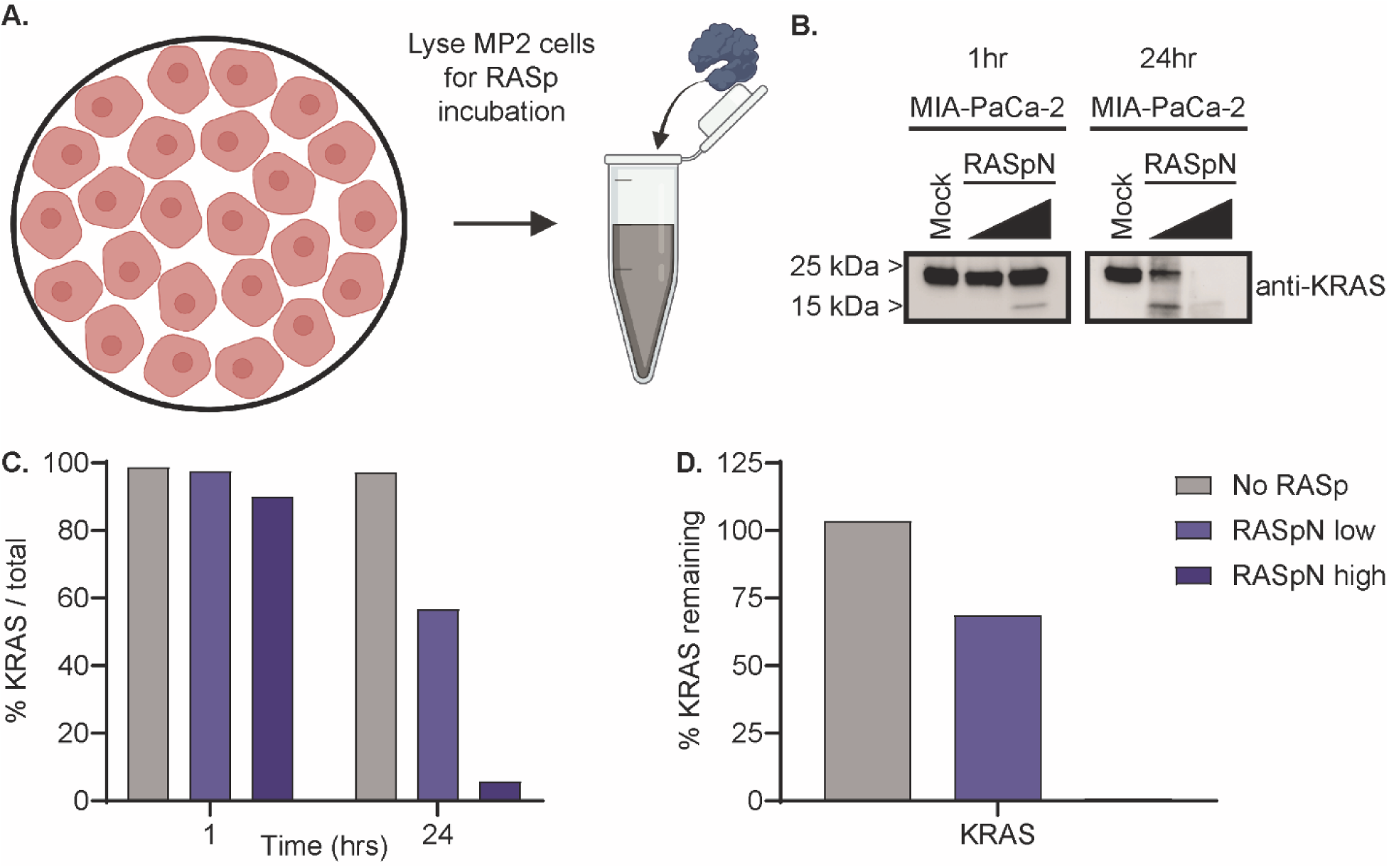
Purified RASp cleaves endogenous KRAS expressed in MIA-PaCa-2 cell lysates. (**A**) Experimental design schematic created with BioRender.com. MIA-PaCa-2 cells were lysed in RIPA buffer prior to incubation with RASp. (**B**) Western blot shows cleavage of MP2 cell lysates expressing KRAS after incubation with RASpN in a dose-dependent manner for 1 hr, left, or 24 hr, right. (**C**) Quantification of KRAS expression signal by KRAS detection in (A) using FIJI. (**D**) Quantification of remaining KRAS expression in (A) expressed as ratio of KRAS signal remaining at 24 hr divided by the KRAS signal at 1 hr using FIJI.

After confirming that RASpN activity in MP2 cell lysates, we transfected MP2 cells with RASpN and induced with doxycycline 24 hpt. MP2 cells were then lysed 72 hpt, as indicated by the cartoon (**Figure 5A**) and assayed by western blot for expression and depletion of KRAS (**Figure 5B**). To further understand these results, we assayed these transfected cells by immunofluorescence microscopy to determine expression of FLAG on a per cell basis and to monitor cell health (**Figure 5C**). We note the low levels of FLAG signal upon transfection and induction of MP2 cells with RASpN. Nonetheless we do observe a significant decrease in total cells, as determined by counting nuclei upon expression and induction of RASpN compared with either mock-transfected or uninduced MP2 cells (**Figure 5D**). Importantly, even with the modest expression of RASpN in MP2 cells upon transfection, we are able to observe substantial cell killing.

**Figure 5.**
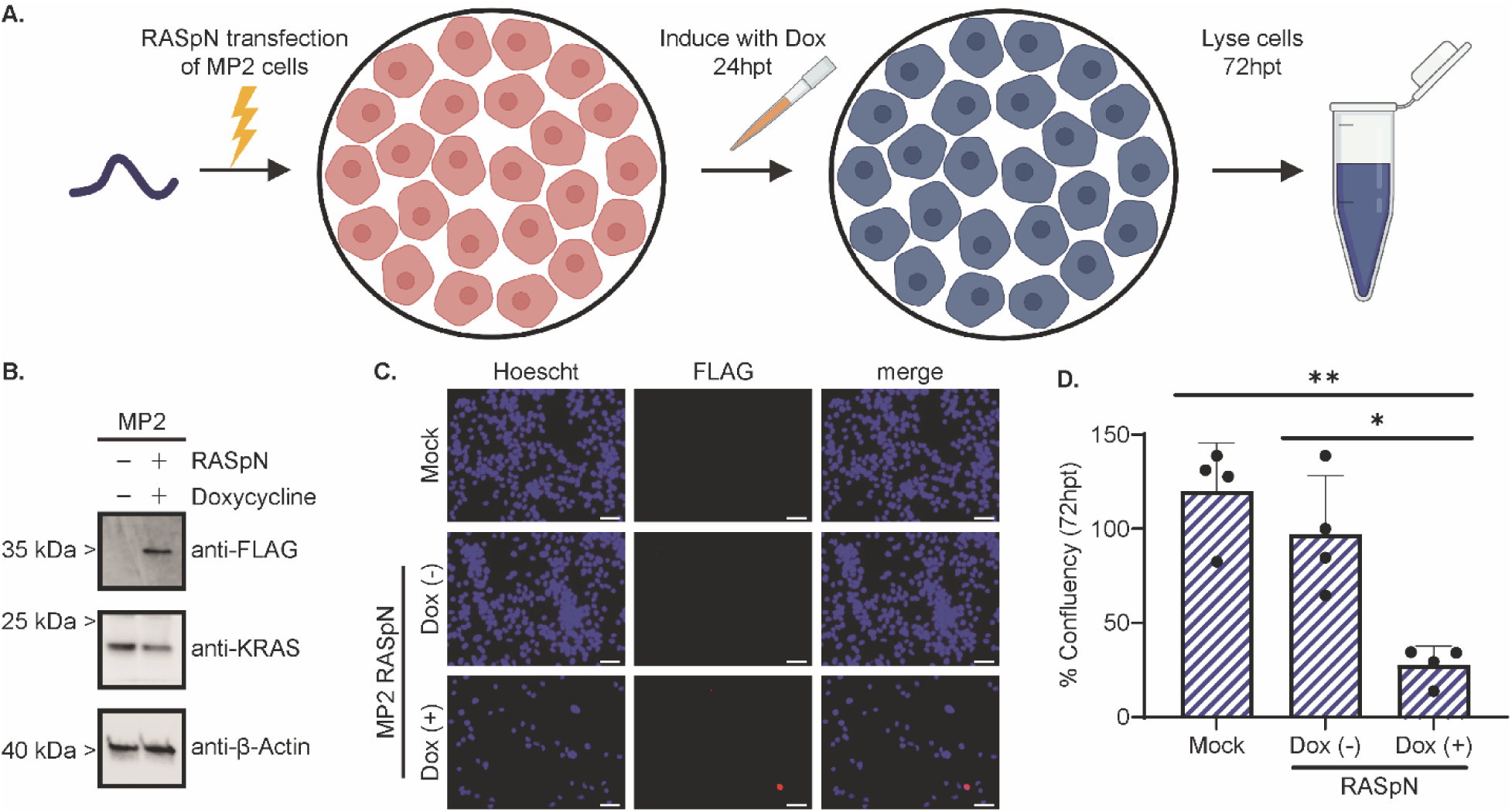
RASpN cleaves endogenous KRAS expressed in MP2 cells upon transfection. (**A**) Experimental design schematic created with BioRender.com. MIA-PaCa-2 cells were transfected with RASpN. 24hpt RASpN was induced with doxycycline, and cells were lysed in RIPA buffer 72hpt. (**B**) Western blot identifies cleavage of KRAS in MP2 lysates after transfection with tetracycline-inducible RASpN or mock-transfected. (**C**) Immunofluorescence-mediated detection of RASpN (FLAG; red) in MP2 cells transfected in parallel with a tetracycline-inducible RASpN or mock-transfected. RASpN-transfected cells were treated with doxycycline 24hpt, with staining performed 72hpt. Nuclei (Hoescht; blue). Scale bar = 100 µm. (**D**) Quantification of % confluency from panel (B). Each bar represents mean assayed across n=4 images taken at predetermined locations within the culture. Error bars represent standard deviation across images. Statistical analysis was performed using a paired t-test (* p <0.05, ** p <0.01).

To further enhance RASpN expression in MP2 cells, we packaged RASpN in a lentiviral delivery vector to stably-transduce our RASpN construct in MP2cells and control HEK 293T cells. Cells were grown under puromycin selection to maintain a pure population of MP2 cells stably-expressing RASpN as indicated (**Figure 6A**). Cell lysates were collected 24 hpi, and MP2 lysates were probed by western blot alongside MP2 cells transfected with either a non-targeting control (NTC) or KRAS-specific siRNA. Importantly, we observe robust expression of RASpN by blotting for FLAG and a strong decrease in total KRAS upon transduction and induction of RASpN or by siRNA (**Figure 6B**). However, most importantly this system provides a platform to understand how RASpN affects cell signaling pathways downstream of KRAS expression (**Figure 6C**). We observe a strong depletion of both MEK and phosphorylated-MEK (pMEK) proteins upon transduction and induction of RASpN, which is not seen using the KRAS-targeting siRNA. Further downstream of MEK, we observe a modest depletion of ERK and pERK upon transduction and induction of RASpN. As part of the mTOR pathway, we also observe a near total ablation of AKT upon expression and induction of RASpN. Zooming in to observe the changes on a per cell basis, we examined MP2 cells stably-transduced with RASpN and upon doxycycline induction at 0, 6, 12, and 24 hpi by immunofluorescence microscopy assaying both RASpN expression by FLAG and for cell health by staining nuclei (**Figure 6D**). We observe peak expression of RASpN 12 hpi, and see a significant decrease in living cells throughout the time course, with only single digit cells remaining 24 hpi (**Figure 6E**). To assess whether RASpN expression differentially affects cells harboring mutant versus WT KRAS, we directly compared the cell killing rates between 293T cells which express low levels of endogenous WT KRAS, and MP2 cells that contain endogenous KRAS-G12C. Thus, upon stable expression, puromycin selection, and doxycycline induction, we observe a marked and significantly more rapid cell killing for MP2 cells compared with 293T cells, 24 hpi (**Figure 6F**). For MP2 cells, RASpN is able to kill nearly all cells by 24 hpi, whereas in that same time frame more than 50% of the 293T cells remain viable (**Figure 6G, H**). This observation provides strong evidence for the selective preference of RASpN killing those cells with KRAS locked in a GTP-bound state over cells with WT KRAS cycling between GTP-bound and GDP-bound states.

**Figure 6.**
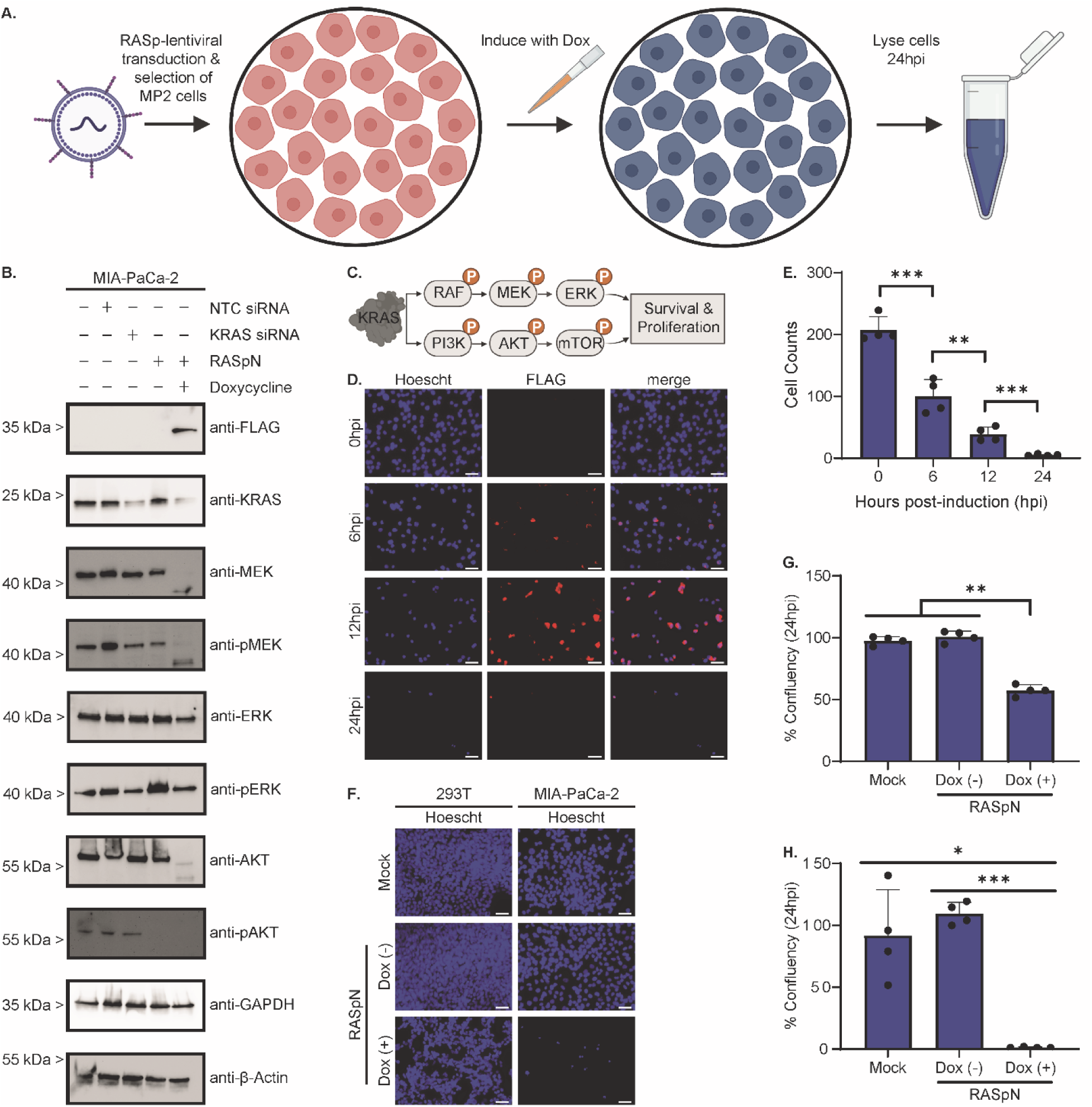
RASpN cleaves endogenous KRAS expressed in MP2 cells upon lentiviral transduction and prevents downstream effector activation and signaling. (**A**) Experimental design schematic created with BioRender.com. MIA-PaCa-2 cells were transduced with a RASpN-expressing tetracycline-inducible lentivirus. was induced with doxycycline, and cells were lysed in RIPA buffer 24hpi (hours post-induction). (**B**) Western blot identifies cleavage of KRAS in MP2 lysates after transduction with tetracycline-inducible lentivirus. MP2 cells were mock-treated or transfected with non-targeting control siRNA or KRAS-specific siRNA as controls. (**C**) KRAS downstream signaling cascade created with BioRender.com (**D**) Immunofluorescence-mediated detection of RASpN (FLAG; red) in MP2 cells transduced with a tetracycline-inducible RASpN. Cells were treated with doxycycline 24hps (hours-post seeding). Cells were fixed at 0, 6, 12 and 24hpi. Nuclei (Hoescht; blue). Scale bar = 100 µm. (**E**) Cell counts from panel (D). Each bar represents mean assayed across n=4 images taken at predetermined locations within the culture. Error bars represent standard deviation across images. Statistical analysis was performed using a paired t-test (** p <0.01, *** p <0.001). (**F**) Immunofluorescence-mediated detection of RASpN (FLAG; red) in 293T, left, MP2, right, transduced with a tetracycline-inducible RASpN. Cells were treated with doxycycline 24hps (hours-post seeding). Cells were fixed 24hpi. Nuclei (Hoescht; blue). Scale bar = 100 µm. (**G**) Quantification of % confluency from 293T cells in panel (F). (**H**) Quantification of % confluency from MP2 cells in panel (F). Each bar represents mean assayed across n=4 images taken at predetermined locations within the culture. Error bars represent standard deviation across images. Statistical analysis was performed using a paired t-test (* p <0.05, ** p <0.01, *** p<0.001).

## Discussion

Targeting active RAS to control its signaling pathways, either as a therapeutic against RAS-driven cancers or as a reagent, is an important goal that has been sought via a variety of approaches that include small molecule inhibitors (*11–23*), PROteolysis TArgeting Chimeras (PROTACs) that stimulate RAS degradation (*24*), monobodies that inhibit RAS signaling (*25*), and proteases that degrade RAS directly (*8, 26–32*). Our previous work (*8*) was focused on developing a protease that site-specifically cleaves RAS at switch II, thereby leveraging an order-to-disorder transition that this region undergoes as it adopts the GTP-bound active state (*7–10*). The ability to control signaling via degradation and depletion of RAS is a key test for our designed RASProtease, so we examined its ability to cleave RAS and regulate downstream signaling pathways in the pancreatic cancer cell line MP2. In these studies, we showed that mutant KRAS was depleted in these cells upon induction of RASProtease expression and that depletion of KRAS coincided with a decrease in signaling via the MEK-ERK pathway. Moreover, depletion of KRAS by RASProtease induced nearly complete cell death in MP2 cells within 24 hours of induction. These studies demonstrate that RASProtease can control RAS signaling pathways and consequently cell health in relevant model systems.

Cofactor activation is a novel means whereby protease activity can be more tightly regulated and here we systematically assessed the effectiveness of the nitrite- and imidazole- dependent versions (*8*) of our RASProtease. We first showed that RASpN cleaves a fluorescently-labeled peptide substrate with a *k_obs_* that is approximately 100-times faster than does RASpI (**Fig. 1**). When tested against intact KRAS, we observe the same phenomenon, regardless of nucleotide or mutational status. KRAS cleavage is observed for both proteases, but RASpI cleavage lags far behind that of RASpN (**Fig. 2C-F**). In each experiment, the cofactor is present at a concentration that is far in excess of the protease concentration and at levels that gave maximal activity in previous studies (*8*). This observation was also replicated in cell lysates (**Fig. 2**) and in the eGFP-KRAS HEK 293T reporter model. RASpI can self-activate to remove the prodomain (**Fig. 3B**, lanes 2, 7, and 8), but little eGFP-KRAS cleavage is evident (**Fig. 3B**, lane 8, **Fig. 3C**). The observed self-activation might be due to the high local concentration of the prodomain cleavage site even though endogenous imidazole concentrations might be very low. It is not clear if the observed minimal activity of RASpI in the eGFP-KRAS reporter system is due to a lack of exogenous imidazole entry into cells, or the lower inherent activity of RASpI. It is likely that, either for use as a potential therapeutic or reagent, the cofactor imidazole will have to be delivered in addition to RASpI as has been done previously (*33*). This could provide an extra element of control, as the cofactor for RASpN is sufficiently abundant that no exogenous nitrite is required (e.g. **Fig. 3B**, lane10).

Another consideration when attempting to control RAS signaling via proteolytic cleavage is that active RAS should be the preferred target. Our previous results (*8*) showed that both RASpN and RASpI preferentially cleave active RAS as approximated by the GMPPNP bound protein. Expanding on those results here, we found that at stoichiometric ratios, both RASpI and RASpN preferentially cleave inactive KRAS. However, when RASpN is limiting (in this case at a ratio of 1:10,000 relative to the KRAS concentration), the cleavage preference switches to active KRAS. We assessed preferential cleavage of active KRAS both by *in vitro* cleavage followed by SDS-PAGE analysis (**Fig. 1C-H**) and by NMR analysis of peak intensity changes (**Fig. 1J-K**). The reason for this switch in cleavage preference as a function of RASProtease concentration is unclear. It is possible that, at stoichiometric ratios, RASpN can influence the conformational equilibrium of switch II in the GDP-bound form more readily than the GMP-PNP bound form, whereas in limiting concentrations RASpN must rely on the inherent conformational exchange in switch II of the two forms of KRAS, which we previously showed (*8*) to be much more pronounced in the GMP-PNP bound form. The limiting protease conditions are more likely to be relevant to studies in cells or therapeutic trials as it is unlikely that stoichiometric amounts of protease could be achieved under any reasonable experimental scenario pertinent to those studies.

A final consideration we explored using RASpN was whether its activity in cells would preferentially affect cells that are RAS-dependent. We chose as our cancer model MP2 cells, which exhibit strong KRAS-G12C dependence for growth and survival and exhibit an apoptotic response when KRAS is silenced by siRNA (*34, 35*). By contrast, control HEK 293T cells are not RAS-dependent (*36, 37*). Both cell lines were transduced with lentivirus and RASpN expression was induced under the same conditions. Appearance of RASpN in cells, as judged by the anti-FLAG detection via fluorescence microscopy (**Fig. 6D**) coincides with a marked decrease in MP2 cell viability, with the majority of the cells dead within 12 hpi and few remaining cells at 24 hpi (**Figs. 6D-H**). For HEK 293T cells, at 24 hpi greater than 50% of the cells were viable (**Fig. 6G** and **F**). This disparity is consistent with the fact that MP2 cells are KRAS-dependent. Moreover, because RASpN cleaves all three RAS isoforms and targets active RAS preferentially, these data indicate that the designed RASpN could be a potential therapeutic against RAS-dependent cancers, similar to the Vibrio vulnificus Ras/Rap1 site-specific endopeptidase RRSP (*26–32*). Further improvements in our designer RASpN approach can focus on increased specificity for active RAS, enhanced activity, and tailoring cofactor dependence based on the unique metabolic profiles of specific cancer cells.

## Materials and Methods

### Plasmids

Tetracycline inducible RASpI and RASpN have previously been described (*8*), while RASp* contains a S221A mutation, rendering it catalytically inactive, in addition to K101S. eGFP-KRAS was synthesized from was a kind gift from Dominic Esposito (The RAS Initiative). VSV-G and psPAX2 were purchased from Addgene.

### Tissue Culture

HEK 293T (293T; ATCC) cells, maintaining wild-type KRAS, and MIA-PaCa-2 (ATCC) cells, expressing KRAS G12C mutant, were grown in DEM with 10% heat-inactivated, tetracycline-free fetal bovine serum (TF-FBS), maintained at 37°C with 5% CO_2_.

### Protein Expression and Purification

Constructs for prodomain-RASpN and prodomain-RASpI were cloned into the pJ1 expression vector (*38*). WT-KRAS was cloned into the eXact tag pH0720 expression vector, which contains both an engineered subtilisin prodomain tag and an HSA-binding GA sequence as N-terminal fusion domains. Both plasmids were transformed into *E. coli* BL21(DE3) cells (New England Biolabs). Protein expression for the protease constructs were conducted in LB media, while WT-KRAS was grown in M9 minimal media for ^15^N labeling, as previously described (*39*). Following cell harvesting and sonication, the protease lysates were cleared by centrifugation and the supernatant fraction was incubated with 10 mM sodium nitrite or 10 mM imidazole for RASpN and RASpI, respectively, at 25 °C for 1 hour to cleave the prodomain. The cleaved samples were subsequently purified by affinity chromatography on a column derivatized to the QEEYSAM peptide, as previously described (*39*). Pure fractions were combined and concentrated for downstream experiments.

WT-KRAS was purified by affinity chromatography on a 5 mL subtilisin column, followed by a 1 mL human serum albumin (HSA) column. The subtilisin column was first equilibrated with 5 column volumes (CV) of room temperature running buffer (100 mM potassium phosphate, pH 7.0). The lysates (6 mL) were loaded onto the equilibrated column and washed as follows: 5 CV of cold running buffer, 20 CV of cold wash buffer (100 mM potassium phosphate, 500 mM sodium chloride, pH 7.2), 5 CV of room temperature running buffer, and finally 2 CV of elution buffer (100 mM potassium phosphate, 3.5 mM imidazole, pH 7.0). The eluent was collected as 1 mL fractions and then transferred to a 10 mL syringe. After equilibration of the HSA column with 8 CV of running buffer, the eluent was flowed through the column to ensure separation of the uncleaved fusion protein from the cleaved WT-KRAS, which was collected in the flow-through. An additional 5 CV of running buffer was flowed through the column to collect any remaining cleaved target protein. SDS-PAGE analysis was conducted to confirm protein purity. The flow-through, containing WT-KRAS in the GDP-bound form, was then concentrated for subsequent experiments.

GMPPNP-bound KRAS was prepared from GDP-bound KRAS using an enzyme-assisted nucleotide exchange method. Purified GDP-bound KRAS was buffer exchanged into 20 mM Tris-HCl, pH 7.5 containing 100 mM sodium chloride and diluted to 50 µM. A reaction mixture containing final concentrations of 1 mM GMPPNP, 5 mM magnesium chloride, 1 mM TCEP, and alkaline phosphatase (10 U/mg KRAS) was added to the KRAS solution for overnight incubation at 4°C. The samples were then buffer exchanged into 20 mM HEPES, pH 7.4 containing 50 mM sodium chloride, 5 mM magnesium chloride, and 1 mM TCEP, and concentrated.

### Stopped-flow Kinetics

Kinetic measurements were conducted using the Applied Photophysics SX20 stopped flow spectrometer using an excitation wavelength of 360 nm with a 400 nm energy filter, based on the intrinsic properties of the amino-4-methylcoumarin (AMC) fluorophore, as in the previous study (*39*). Immediately before the experiment, the QEEYSAM-AMC peptide was diluted to the desired concentration in 100 mM potassium phosphate pH 7.0 buffer containing 10 mM sodium nitrite for RASpN or 10 mM imidazole for RASpI. The protease samples were prepared in the corresponding buffer.

The protease and peptide samples were prepared and loaded separately into 3 mL syringes and rapidly mixed at a 1:1 volumetric ratio by the stopped-flow instrument at room temperature to make the final solutions at desired concentrations (100 nM for the QEEYSAM-AMC substrate and 4 µM for the protease). Data were collected until reaction completion, which occurred within 5 seconds for RASpN and within 1800 seconds for RASpI. Three replicate measurement scans were collected for each sample, with each scan generating 10,000 data points.

Experiments were conducted under single-turnover conditions and fluorescence signals were processed as previously described (*39*). Fluorescence progress curves for RASpN and RASpI were fitted by non-linear regression to Equation (1) or Equation (2), respectively, using MATLAB (The MathWorks, Inc. (2024). *MATLAB (Version 24.2.0, R2024b)*. Natick, MA: The MathWorks, Inc.). The resulting *k*_obs_ values were averaged to report the mean ± 1 standard deviation.

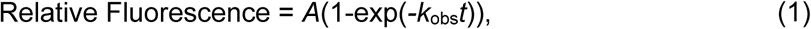

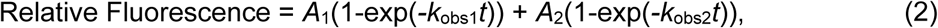

where *A, A*_1_, and *A*_2_ are the amplitudes, *k*_obs_, *k*_obs1_, and *k*_obs2_ are the rate constants, and *t* is the reaction time.

### In Vitro Proteolysis

To assess ability of RASp to cleave its target substrate, KRAS, stoichiometric ratios of purified RASpI or RASpN were incubated with GMPPNP- or GDP-bound WT or G12C KRAS in the presence of necessary cofactors, 10 mM imidazole or 1 mM nitrite, for a fixed duration. In a separate experiment, RASpN was incubated at a 1:10,000 ratio to either wild-type KRAS or G12C KRAS. At the desired timepoint, the proteolysis reaction was stopped by adding SDS buffer containing β-mercaptoethanol and heat inactivated.

### NMR Spectroscopy

All NMR samples were prepared in 95% (v/v) H_2_O/5 % (v/v) D_2_O with a final concentration of 300 µM. A stock solution of RASpN was added at final concentrations corresponding to a stoichiometric (1:1) and a 10,000:1 (KRAS WT: RASpN) ratio. The final buffer conditions for the NMR samples were 0.1 M Phosphate buffer at pH 7.0 with 1 mM nitrite. All NMR experiments were recorded at 37 °C using a Bruker 600 MHz spectrometer with a TCI triple-resonance inverse cryogenic probe featuring x, y, and z gradients. All NMR data were processed using Topspin 4.5.0. Proteolytic digestion of ^15^N-labeled KRAS WT (in GDP of GMP form) in the presence of RASpN (stoichiometric or 10000:1 ratio) was monitored by serially acquired two-dimensional (2D) ^1^H-^15^N TROSY-HSQC experiments in the span of 24 hrs. (for stoichiometric ratio RASpN) or 48 hrs. (for 10000:1 ratio RASpN). Each spectrum was acquired using the same NMR settings (e.g., ns = 32). The total measurement time for each 2D ^1^H-^15^N TROSY HSQC spectrum was 1 hr. and 20 min. ^1^H-^15^N cross-peak intensities (peak height) from each 2D ^1^H-^15^N TROSY-HSQC spectrum were plotted to monitor the proteolytic digestion kinetics.

### Transfection

To assess expression of RASp in 293T, 293T or MIA-PaCa-2 cells (1.0 x 10^5^ cells/well) were seeded in 24-well plates in DMEM (Gibco) with 10% TF-FBS and incubated overnight at 37 °C / 5% CO_2_. The next day, cells were transfected using JetPrime (#1114-07, Polyplus) according the manufacturer’s protocol with the exception of using double the amount of plasmid DNA and JetPrime reagent. At 18 hours post-transfection (hpt), the transfection-containing media was aspirated and replaced with fresh 10% TF-FBS-containing DMEM. At 24 hpt, RASp was induced with 20 µg/mL doxycycline. At 72hpt (48 hours post-induction, hpi), the cells were collected in RIPA buffer containing 2X protease/phosphatase inhibitors.

For downstream immunofluorescence assays, cells were seeded in 8-well chamber slides (#CCS-8, MatTek Corporation) with 5.0 x 10^4^ cells/well and transfected as done for 24-well plates with all volumes reduced by 50%. At indicated experimental timepoints, cells were fixed with 4% paraformaldehyde for 15 min.

### Generation of Lentivirus Stocks

6-well plates were coated with Poly-L-lysine (3P8920, Sigma-Aldrich), washed with cell-grade H_2_O, and allowed to dry prior to seeding HEK 293T cells (4.0 x 10^5^ cells/well). Cells were grown overnight at 37 °C / 5% CO_2_ in Dulbecco’s Modified Eagle Medium (DMEM) containing 10% FBS to achieve 80% confluency the next day. Cells were then washed with PBS and transfected with psPAX2 (Addgene plasmid #12260), pCMV-VSV-G(Addgene plasmid #8454) and RASpN expression plasmid using JetPrime (Polyplus) according to the manufacturer’s protocol 1 µg of RASpN plasmid alongside 200 ng VSV-G glycoprotein, and 700 ng PsPax2 packaging plasmid per well. At 6 hpt, media was replaced with DMEM / 3% FBS and lentivirus-containing supernatant was collected at 72 hpt. Harvested supernatants were pooled, clarified by centrifugation (2000 x g, 10 min, 4° C) and filtered (0.2 micron). HEPES (#15630-080, Gibco Laboratories) and polybrene (#TR1003G, Fisher Scientific) were added to achieve final concentrations of 20 mM and 4 µg/mL, respectively, and lentiviral stocks were stored at −80 °C prior to use.

HEK 293T and MIA-PaCa-2 cells were seeded into 12 well plates at 7x104 cells/well and incubated overnight at 37 °C. The next day, cells were transduced with 400 µL of lentivirus stock in a final volume of 2 mL DMEM / 10% FBS containing 20 mM HEPES and 4 µg/mL polybrene via centrifugation (1000 x g) for 1 hr at 37° C). Following spinoculation, cells were returned to the incubator for 6-18 hrs at which point the media was replaced with 1 mL 185 DMEM / 3% FBS until the transduced phenotype was observed and / or cells were required to be split. hCDHR3-containing cells were selected using 5 µg/mL Puromycin.

### SDS-PAGE and Western Blot

Cell lysates were collected in RIPA buffer containing protease and phosphatase inhibitors and total protein was quantified using a Pierce BCA assay (#23227, Thermo Fisher Scientific). Protein lysates (20 µg) were denatured by heating (98 °C, 2min) and separated on Bio-Rad 4-15% Mini-PROTEAN TGX Stain-Free Protein Gels (#4568085; Bio-Rad) or 4-20% Mini-PROTEAN TGX Stain-Free Protein Gels (#4561095; Bio-Rad) under reducing conditions. Proteins were transferred to PVDF membranes (Bio-Rad). Membranes were blocked in 5% milk diluted in tris-buffered saline with 0.1% Tween-20 (TBS-T) for 1 hour at room temperature with rocking. Primary antibodies were diluted in blocking buffer per Table X. Following overnight incubation with rocking at 4 °C membranes were washed 3 times for 15 minutes each with TBS-T before incubation with HRP-conjugated donkey anti-Rabbit IgG (1:10,000) or HRP-conjugated goat anti-mouse IgG (ab205719; Abcam). Anti-β-Actin-HRP conjugated antibody (#A3854, Sigma-Aldrich) was diluted 1:35000 in 5% TBS-T and incubated at room temperature for one hour. Western blots were visualized with Clarity Western ECL substrate (1705061; Bio-Rad) and imaged using ChemiDoc^TM^ Imaging System (Bio-Rad).

### Immunofluorescence

Following fixation, cells were washed 3 times with PBS and then permeabilized with 0.2% Triton X-100 in PBS for 15 min. After permeabilization, cells were blocked with 3% BSA in PBS for 1 hr at room temperature. Anti-FLAG antibody was diluted 1:200 in 1% BSA / PBS. Following overnight incubation at 4 °C, cells were washing with PBS 3 times prior to secondary antibody staining. Secondary antibodies, AlexaFluor555 anti-mouse IgG 1:500 was applied for 1 hour at room temperature in a humidified chamber. Cells were washed with PBS and incubated with Hoescht (1:2000, 10 min; Thermo Fisher Scientific). Images were acquired with a Zeiss Observer Z1 inverted fluorescence microscope.

## Acknowledgements

We thank Dr. Jaekyun Jeon, Institute for Bioscience and Biotechnology Research (IBBR), University of Maryland, for providing access to the stopped-flow instrument used in the kinetics experiments.

## Author contributions

M.E.G, B.C., K.J.C., J.O., E.A.T., and T.R.F. designed the experiments and analyzed the data. M.E.G, B.C., and K.J.C. performed the experiments. M.E.G., E.A.T., and T.R.F. wrote the manuscript. M.E.G., J.O., E.A.T., and T.R.F. edited the manuscript.

## Data and materials availability

All data required to evaluate the conclusions presented here are in the paper.

---

**Table 1.** Quantification of western blot intensity in Fig 6B; normalized to RASpN(-Dox).

|  | Mock | NTC siRNA | KRAS siRNA | RASpN (-Dox) | RASpN (+Dox) |
| --- | --- | --- | --- | --- | --- |
| KRAS | 126 | 103 | 32 | 100 | 19 |
| B-Actin | 102 | 147 | 138 | 100 | 96 |
| MEK | 171 | 162 | 157 | 100 | 9 |
| pMEK | 218 | 312 | 142 | 100 | 27 |
| ERK | 103 | 98 | 97 | 100 | 64 |
| pERK | 41 | 59 | 38 | 100 | 53 |
| AKT | 125 | 87 | 125 | 100 | 10 |
| pAKT | 986 | 910 | 667 | 100 | 165 |

